# Lipid-polymer hybrid nanoparticles utilize B cells and dendritic cells to elicit distinct antigen-specific CD4^+^ and CD8^+^ T cell responses

**DOI:** 10.1101/2022.01.23.477398

**Authors:** Michael H. Zhang, Brianna L. Scotland, Yun Jiao, Emily M. Slaby, Nhu Truong, Georgina Stephanie, Ryan M. Pearson, Gregory L. Szeto

## Abstract

Antigen presenting cells (APCs) have been extensively studied for treating cancers and autoimmune diseases. Dendritic cells (DCs) are potent APCs that uptake and present antigens (Ags) to activate immunity or tolerance. Despite their active use in cellular immunotherapies, DCs face several challenges that hinder clinical translation, such as inability to control Ag dosing for tuning immune responses and low abundance in peripheral blood. B cells are a potential alternative to DCs, but their poor non-specific Ag uptake capabilities compromise controllable priming of T cells. We developed phospholipid-conjugated Ags (L-Ags) and lipid-polymer hybrid nanoparticles (L/P-Ag NPs) as Ag delivery platforms to expand the range of accessible APCs for use in priming CD4^+^ and CD8^+^ T cells. These delivery platforms were evaluated using DCs, CD40-activated B cells, and resting B cells as a diverse set of APCs to understand the impact of various Ag delivery mechanisms for generation of Ag-specific T cell responses. L-Ag delivery (termed depoting) of MHC class I and II-restricted Ags successfully loaded all APC types in a tunable manner and primed both Ag-specific CD8^+^ and CD4^+^ T cells, respectively. Incorporating L-Ags and polymer-conjugated Ags (P-Ag) into NPs can direct Ags to different uptake pathways to engineer the dynamics of presentation and shape T cell responses. DCs were capable of processing and presenting Ag delivered from both L- and P-Ag NPs yet B cells could only utilize Ag delivered from L-Ag NPs. Multivariate analysis of cytokines secreted from APC:T cell co-cultures indicated that L-Ag NPs primed different T cell responses than P-Ag NPs. Altogether, we show that L-Ags and P-Ags can be rationally paired within a single NP to leverage distinct delivery mechanisms to access multiple Ag processing pathways in two APC types, offering a modular delivery platform for engineering immunotherapies.

## Introduction

Antigen (Ag)-specific immunotherapies can train the host immune system to recognize disease-associated Ags to initiate cellular and humoral immune responses. Despite successes preventing infectious diseases, few Ag-specific immunotherapies or therapeutic vaccines have fulfilled clinical expectations or have been FDA approved as treatments for chronic illnesses such as autoimmune diseases or cancers [1][2][3]. One notable clinical challenge is effectively delivering Ags *in vivo* or *ex vivo* to antigen-presenting cells (APCs), which are the primary mediators of cellular immune responses toward Ag-specific immunity or tolerance. Dendritic cells (DCs) are professional APCs that uptake, process, and present Ags in a MHC class I- and class II-restricted manner to prime CD8^+^ and CD4^+^ T cells, respectively. Leveraging DCs *in vivo* is clinically challenging due to their heterogeneous Ag processing and T cell priming capabilities [4][5]. In addition, their scarcity in peripheral blood, poor proliferation potential, and complex conditioning regimens hinder access for large-scale *ex vivo* engineering. B cells are a promising alternative to DCs due to their high abundance in the peripheral blood and robust expansion *ex vivo* [6][7][8]. These attributes are advantageous for engineering large quantities of APCs to satisfy infusion prerequisites. A major barrier for engineering B cell therapies is their inefficient non-specific uptake of Ags. B cells typically use their B-cell receptors to uptake and process specific Ags, but this capability is contingent on their activation state, a feature of adaptive immunity that is advantageous for promoting specific humoral immunity but precludes non-specific Ag uptake and broad T cell stimulation [9][10][11]. Developing Ag delivery strategies that are mechanistically independent of cellular activation state can leverage existing capabilities of APC candidates and expand their potential to elicit Ag-specific immunity or tolerance to treat chronic diseases.

Engineered Ag delivery strategies can enhance existing Ag-specific immunotherapies by increasing control over Ag dosing, Ag presentation, and Ag-specific T cell responses. For example, currently available methods to promote Ag cross-presentation to activate CD8^+^ T cells include Ag cargo delivery by electroporation, transfection, viral vector-based transduction, or mechanical perturbation using microfluidic systems [12][13][14][15][16][17]. These methods risk damaging membrane integrity and reducing cell viability. Biomaterial-based carriers have been developed for carrying diverse peptide and protein Ags with decreased cellular toxicities and increased cargo protection against enzymatic degradation. Biomaterials comprised of lipids [18][19][20], biodegradable polymers such as poly(lactic-co-glycolic acid) (PLGA), and other polymers [21][22][23][24][25] can form Ag-carrying nanoparticles (NPs) with increased stability, preferred biodistribution profiles, and accumulate within phagocytic APCs when delivered *in vivo*. The direct conjugation of Ags to these biomaterials promotes delivery for increased uptake, MHC-restricted Ag presentation, and Ag-specific T cell activation and proliferation. PLGA-based conjugates are generally insoluble in aqueous environments, and thus require either complexation with or encapsulation in biomaterial-based carriers to promote delivery to APCs. Lipid-based conjugates can promote delivery to a multitude of cell types by inserting into cell plasma membranes for surface presentation and internalization [26][27][28][29][30], mimicking the natural cell-membrane insertion of lipid-anchored proteins and bypassing the need for active uptake by endocytic mechanisms. Lipid-conjugated Ags and adjuvants have also demonstrated therapeutic efficacy *in vivo* by accumulating in lymphoid organs through albumin binding, promoting delivery to lymphoid APCs [31][32][33][34]. A notable disadvantage of some lipid-based conjugates is their low stability *in vivo* [35]. Formulating lipid and PLGA hybrid NPs that strategically combine their physicochemical features can overcome their respective disadvantages and promote Ag and therapeutic vaccine delivery to APCs [36][37][38]. Hybrid NPs offer a versatile, broadly applicable platform technology to engineer Ag-specific T cell responses for immunotherapy.

Here, we describe the development and evaluation of lipid- and PLGA-based Ag delivery systems as a multimodal approach to investigate cell type-dependent induction of Ag-specific T cell responses using DCs, CD40 pathway-activated B cells (CD40 B-APCs), and non-activated B cells (B-APCs), two APC types that have different endocytic capabilities and embody distinct cellular activation states. We conjugated exact MHC I- or MHC II-restricted peptide Ags to lipids (L-Ag) to allow direct insertion into plasma membranes of DCs and B cells. We hypothesized that incorporating L-Ag and PLGA-conjugated Ag (P-Ag) into PLGA NPs, forming lipid-polymer hybrid NPs (L/P-Ag NPs), would enable two distinct mechanisms of uptake to be leveraged in a single platform: (1) endocytosis of PLGA NP for delivery of P-Ag and (2) depoting by L-Ag. Using a series of co-culture experiments and immunoassays with DCs, CD40 B-APCs, or B-APCs, we examined the APC-dependent efficiency of induction of Ag-specific CD4^+^ and CD8^+^ T cell activation, proliferation, and cytokine secretions as a function of L-Ag- or P-Ag-mediated PLGA NP delivery to APCs. We show that L-Ag NPs can take advantages of lipid- and PLGA-mediated NP delivery strategies to engineer APCs to differentially prime CD4^+^ and CD8^+^ T cells. Our findings demonstrate that multimodal mechanisms of Ag delivery can be achieved through the choice of biomaterial-Ag conjugation strategy and that incorporation into NPs does not hinder APC accessibility nor the ability to induce of Ag-specific T cell responses. Our findings have broad therapeutic potential to guide the future design of novel APC-targeted therapeutic interventions for *ex vivo* or *in vivo* Ag-specific immunotherapy applications.

## Materials and Methods

### Materials

Acid-terminated 50:50 poly(lactic-co-glycolic acid) (PLGA) (~0.17 dL/g inherent viscosity in hexafluoro-2-propanol; approximately MW 4.2 kDa) was purchased from Lactel Absorbable Polymers (Birmingham, AL). Poly(ethylene-alt-maleic anhydride) (PEMA; MW 400 kDa) was received as a gift from Vertellus™ (Indianapolis, IN). 1,2-distearoyl-sn-glycero-3-phosphoethanolamine (DSPE)-(poly(ethylene glycol)) (PEG)-2000 N-hydroxysuccinimide lipid (DSPE-PEG2000-NHS) was purchased from Nanocs, Inc (Natick, MA). Amine-terminated Ags OVA257-264 (SIINFEKL), OVA_323-339_ (ISQAVHAAHAEINEAGR), and GP100 (CAVGALEGPRNQDWLGVPRQL) [39] were purchased from GenScript Biotech (Piscataway, NJ). Fluorescein-labeled SIINFEKL was purchased from Anaspec. All other chemicals were purchased from MilliporeSigma (Saint Louis, MO) unless stated otherwise.

### Lipid- and PLGA-Ag synthesis

Lipid-Ag (L-Ag) and PLGA-Ag (P-Ag) conjugates were synthesized by adapting a previous protocol [24]. Peptides Ags were dissolved in a solution of dimethyl sulfoxide (DMSO) at 10 mg/mL. For both L-Ag and P-Ag synthesis, triethylamine (5x molar excess to peptide) was added to the peptide solution under stirring. PLGA was dissolved in DMSO at 20 mg/mL. N-(3-dimethylaminopropyl)-N’-ethylcarbodiimide hydrochloride (EDC) crosslinker was dissolved at 20 mg/mL in DMSO and added dropwise (5x molar excess to peptide) to stirring PLGA solution. N-hydroxysuccinimide (NHS) was dissolved at 5 mg/mL in DMSO and added dropwise (5x molar excess to peptide) to the stirring solution. After 15 min at 25°C, peptide Ag was added dropwise (1.1x molar excess to PLGA) to the stirring NHS-functionalized PLGA solution, and the reaction proceeded overnight at 25°C. For coupling lipids to peptide Ags, DSPE-PEG-NHS was dissolved at 20 mg/mL in DMSO under stirring. The peptide (1.1x molar excess to DSPE-PEG-NHS) was then added drop-wise to the stirring solution, and the reaction proceeded overnight at 25°C. The resulting conjugates were purified by dialysis using 3,500 molecular weight cut-off membrane against 4L distilled water, with six water exchanges over 2 days, and then lyophilized. The coupling efficiency was assessed using ^1^H-NMR analysis and determined as previously described by our group [24]. Fluorescently labeled lipid-GP100 was synthesized as previously described [32] and provided by the Irvine Lab from MIT.

### Nanoparticle production and characterization

PLGA NPs containing L-Ag or P-Ag conjugates were produced by adapting a single emulsion-solvent evaporation method using PEMA as an emulsion stabilizer [24]. PLGA was dissolved at 50 mg/mL in dichloromethane, and conjugates were dissolved at 20 mg/mL in DMSO. L-Ag or P-Ag conjugates were mixed with the PLGA polymer to achieve a final ratio of 8 μg peptide Ag per mg of PLGA. To this, 10 mL of 1% PEMA was added and the mixture was sonicated at 100% amplitude for 30 s using a Cole-Parmer 500-Watt Ultrasonic Homogenizer. The resulting oil-in-water emulsion was then poured into 40 mL of magnetically stirred 0.5% PEMA. The combination of Ags and biomaterials resulted in 4 NP formulations: (1) (L-1/L-2) NP; (2) (L-1/P-2) NP; (3) (L-2/P-1) NP; or (4) (P-1/P-2) NP (**Table 1 and Table 2**). After overnight stirring at 25°C to evaporate dichloromethane, NPs were washed 3 times at 11,000 x *g* for 20 min at 4°C with water and lyophilized overnight with 4% (w/v) sucrose and 3% (w/v) D-mannitol as cryoprotectants. NP size and zeta potential were evaluated using a Zetasizer Nano ZSP (Malvern Instruments, United Kingdom). NPs were reconstituted in water and washed 3 times by centrifugation before use.

**Table 1.** SIINFEKL and OVA_323-339_ Ag coupling efficiencies to DSPE-PEG (lipid) or PLGA biomaterials to form Ag conjugates.

| Biomaterials | Antigen (Ag) | Conjugates | Coupling efficiency | Naming scheme |
| --- | --- | --- | --- | --- |
| DSPE-PEG | SIINFEKL<br>or<br>OVA323-339 | DSPE-PEG-SIINFEKL | 80.5% | L-1 |
|  |  | DSPE-PEG-OVA<sub>323-339</sub> | 76% | L-2 |
| PLGA |  | PLGA-SIINFEKL | 91% | P-1 |
|  |  | PLGA-OVA<sub>323-339</sub> | 68.3% | P-2 |

**Table 2.** Size, polydispersity index (PDI), and zeta potential of NP variations prepared in this study.

| L/P-Ag NP | Size (nm) | Zeta Potential (mV) | PDI |
| --- | --- | --- | --- |
| (L-1/L-2) NP | 734.3 ± 23.4 | -29.8 ± 0.2 | 0.397 |
| (L-1/P-2) NP | 811.3 ± 49.2 | -29.4 ± 0.5 | 0.273 |
| (L-2/P-1) NP | 763.1 ± 14.7 | -33 ± 2.4 | 0.143 |
| (P-1/P-2) NP | 709.7 ± 33.5 | -33.1 ± 0.3 | 0.38 |

### Isolation of mouse immune cells

All procedures with animals and animal-derived materials were approved by the UMBC Institutional Animal Care and Use Committee (OLAW Animal Welfare Assurance D16-00462). C57BL/6 mice, OT-I (C57BL/6-Tg(TcraTcrb)1100Mjb/J), and OT-II (B6.Cg-Tg(TcraTcrb)425Cbn/J) transgenic mice from Jackson Laboratory (Bar Harbor, ME) were bred in the UMBC animal facility for experimental use. For OT-I and OT-II T cells, spleens from respective OT-I and OT-II mice (<6 months old) were mashed through a 40 μm cell strainer treated with ACK lysing buffer (1 mL per spleen, ThermoFisher Scientific; Waltham, MA) for 5 min at 25°C to lyse red blood cells. OT-I T cells were isolated using a negative selection cocktail containing the following biotinylated mouse antibodies (BioLegend; San Diego, CA): TCR γ/δ (clone GL3), CD24 (clone M1/69), TER-119 (clone TER-119), CD49b (clone HMα2), CD45R/B220 (clone RA3-6B2), CD19 (clone 6D5), CD11c (clone N418), CD11b (clone M1/70), and CD4 (clone: H129.19). OT-II T cells were isolated using the same cocktail, except the CD4 antibody was exchanged with a biotinylated CD8 antibody (clone: 53-6.7). B cells were isolated from spleens of naïve C57BL/6 mice and used the same antibody cocktail, except both CD4 and CD8 antibodies were included while CD45R/B220 (clone RA3-6B2) antibody was removed. Antibody-bound cells were depleted with streptavidin RapidSphere™ magnetic beads according to the manufacturer’s instructions (STEMCELL Technologies; Vancouver, Canada). Cells with >90% purity were used for experiments. CD40-activated B cells (CD40 B cells were acquired by suspending isolated B cells at 2 x 10^6^ cells/mL in complete RPMI 1640 media supplemented with 10% fetal bovine serum (FBS) (ThermoFisher Scientific), and stimulated with 1 μg/mL R848 (TLR7/8 agonist; InvivoGen, Inc.; San Diego, CA) and 5 μg/mL of CD40 antibody ligand (clone: HM40-3; BioLegend).

Bone marrow-derived dendritic cells (BMDCs) were generated from bone marrow of C57BL/6 mice by culturing for 9-11 days with complete RPMI 1640 media supplemented with 10% FBS, 50 mM of β-mercaptoethanol, 1% Pen-Strep, and 20 ng/mL of granulocyte-macrophage colony-stimulating factor (GM-CSF) (Peprotech, Inc.; Rocky Hill, NJ). Supplemented RPMI media was exchanged on days 3, 6, and 8 as previously described [24].

### Ag delivery and co-culture assays

B cells, CD40 B cells, and BMDCs were pulsed with exact Ags or depoted with L-Ags at 10 μg/mL for 1 h in complete RPMI media in a 96-well round bottom plate or 1.5-mL Eppendorf tubes. APCs that were not pulsed or depoted instead received 10 μg/mL of Ags in solution, or loaded at 150 μg/mL for 1 h with various NP formulations. All loaded APCs were washed 3 times in PBS supplemented with 1% bovine serum albumin (BSA) (MilliporeSigma). OT-I and OT-II cells were labeled with 5 μM of carboxyfluorescein succinimidyl ester (CFSE, ThermoFisher Scientific). Ag-loaded APCs were cultured with equivalent number of OT-I cells, OT-II cells, or both T cells (20,000 of each cell type). After 3 days, cells were incubated with αCD16/32 antibody (clone 93; BioLegend) to block non-specific antibody binding by Fc receptors for 5 min at 25°C, before staining with CD4 (clone GK1.5; PerCP/Cyanine5.5), CD8a (clone 53-6.7; APC), and CD25 (clone PC61; PE). Cell proliferation was measured by CFSE dilution. Cell proliferation index and division index were calculated using FlowJo LLC software. Proliferation index is defined as the total number of cell divisions divided by the number of divided cells, whereas division index is the total number of cell divisions divided by the number of total original cells.

Supernatants from co-cultures were collected on day 3 and analyzed using Luminex bead-based multiplex assay (MILLIPLEX MAP Mouse Cytokine/Chemokine Magnetic Bead Panel; MilliporeSigma, Burlington, MA), using manufacturer’s instructions with the following modifications: magnetic capture beads and detection antibodies were used at 0.2x the recommended concentrations.

### Statistical and Multivariate Analyses

All statistical analyses were performed with Prism software (GraphPad Software; San Diego, CA). Multivariate analyses were performed using the “mixOmics” package in R version 4.1.0 [40].

## Results

The ability of lipids to insert into plasma membranes can be exploited for biomimetic delivery of lipid-conjugated Ags into the membranes of cells. We previously demonstrated depoting of lipid-conjugated TLR ligands into immune cell surfaces and endosomes to adjuvant immune responses [30]. Here, we extend the concept of depoting to deliver Ags to APCs for promoting processing and MHC-restricted presentation, both as free conjugates and encapsulated in polymeric NPs (**Figure 1**). We conjugated MHC-restricted peptide Ags derived from protein ovalbumin, SIINFEKL and OVA_323-339_, to DSPE-PEG-NHS using carbodiimide chemistry to yield DSPE-PEG-Ag conjugates (L-Ag) (**Figures 1A and S1**). Coupling efficiencies were determined by ^1^H-NMR characterization for DSPE-PEG-SIINFEKL (80.5%) and DSPE-PEG-OVA__323-339__ (76%) (**Figure S1 and Table 1**). Fluorescently labeled L-SIINFEKL-FAM depoted into bulk splenocytes with high efficiency, including non-endocytic cell types (B220^+^ B cells (B-APCs), CD3^+^ T cells) and endocytic CD11c^+^F4/80^−^ DCs (**Figure 2A-D**). Additionally, the loading of L-Ag was increased compared to Ag only control in B-APCs and T cells by 1.7- and 1.5-fold, respectively (**Figure 2B, C**). No loading increase was observed in DCs with L-Ag due to their high endocytic capabilities (**Figure 2D**).

**Figure 1.**
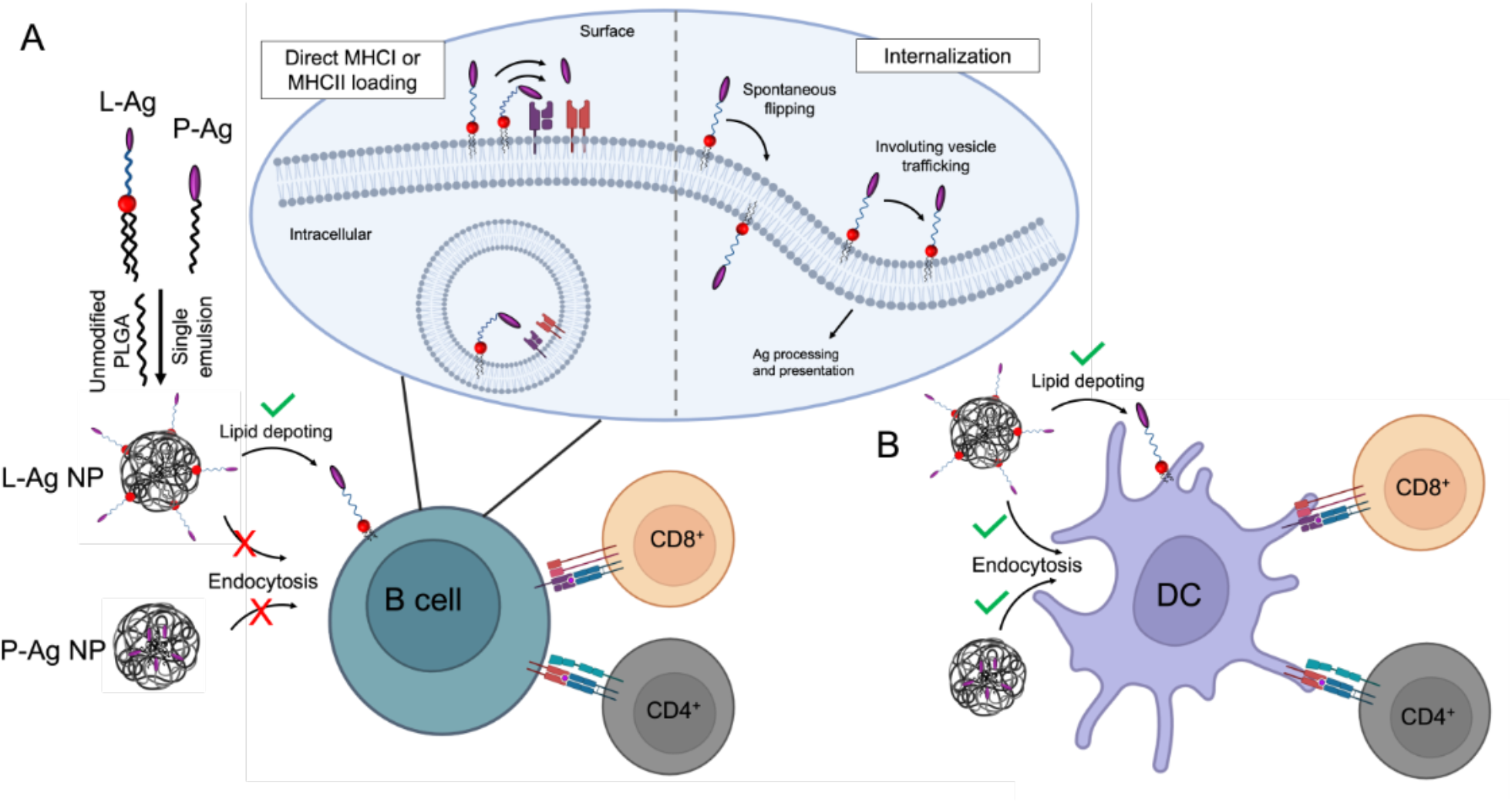
Schematic of lipid-Ag (L-Ag) and PLGA-Ag (P-Ag) NP formulation and proposed mechanisms of Ag delivery to B cells and dendritic cells (DCs). **A)** Ags are conjugated to the end group of lipid or PLGA, combined with unmodified PLGA, and formulated into L-Ag or P-Ag NPs for delivery to APCs. Depoting of L-Ag into the B cell plasma membrane allows an additional mechanism of drug delivery to B-APCs through possible mechanisms, including loading of exact peptide epitopes to surface MHCs, spontaneous flipping of L-Ag into cell cytosol, and through endosomal trafficking to enable Ag presentation for CD4^+^ or CD8^+^ T cell activation. **B)** Depoting and NP-directed endocytosis can access DCs for Ag delivery, processing, and MHC-restricted presentation to activate CD4^+^ or CD8^+^ T cells.

**Figure 2.**
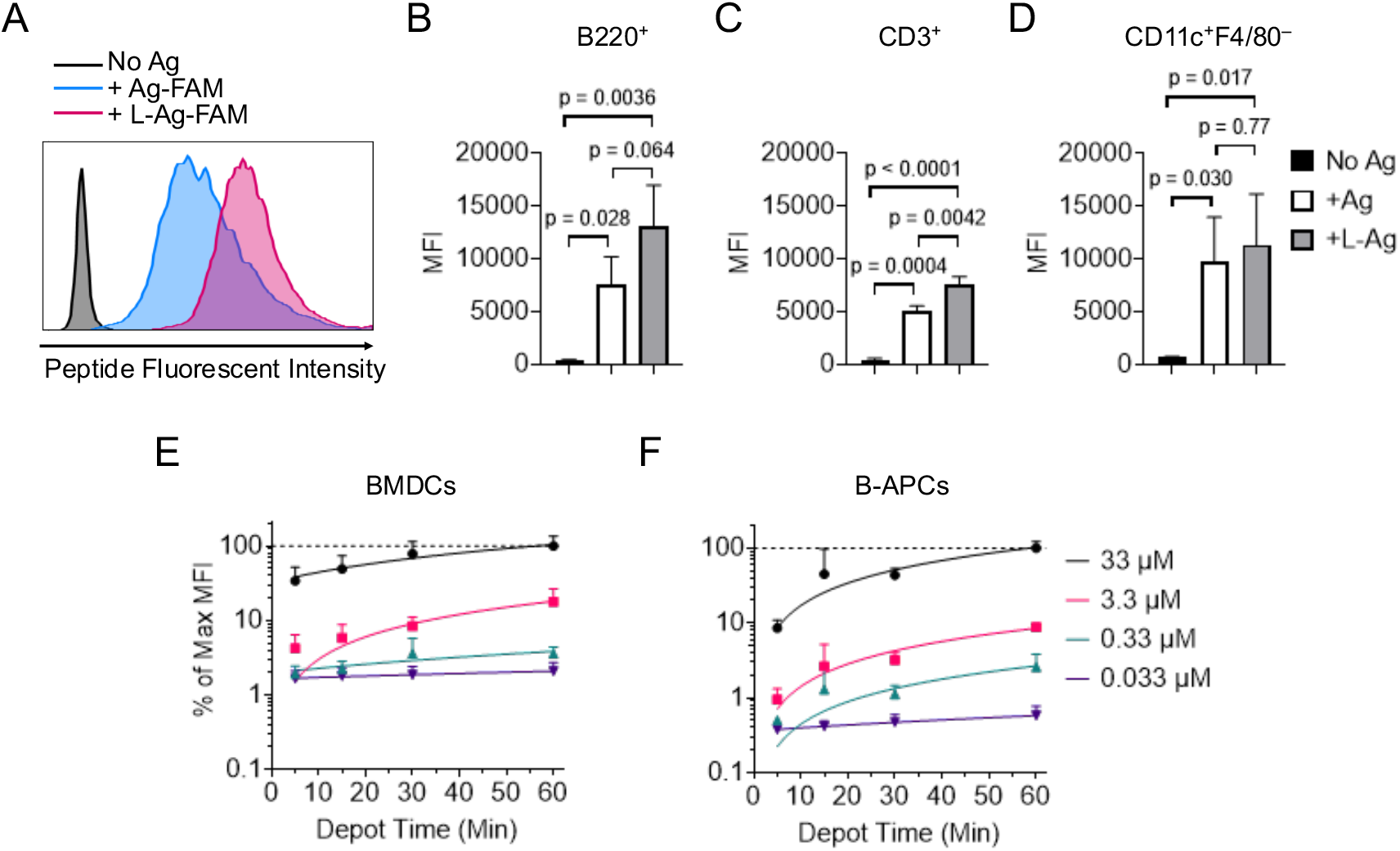
Lipid domain enhances Ag delivery to immune cells and provides tunable dose control. **A)** Fluorescein (FAM) labeled SIINFEKL antigen (Ag) or lipid-conjugated SIINFEKL peptide (L-Ag) was incubated with splenic immune cells at 10 μg/mL for 1 hour 37°C in supplemented RPMI media. Representative fluorescent intensity as measured by flow cytometry. Quantification of fluorescent intensity shown for **B)** B220^+^ cells, **C)** CD3^+^ cells, **D)** CD11c^+^F4/80^−^ cells. Data shown as mean ± s.d. n = 3 independent samples. p values between indicated conditions were determined by RM one-way ANOVA with Tukey’s method for multiple comparisons correction. **E–F)** Lipid-conjugated 20-mer peptide Ag, GP100 (CAVGALEGPRNQDWLGVPRQL) with fluorescein label was loaded on BMDCs and B-APCs. Relative density of L-GP100 loading is dependent on loading concentration and time, as analyzed by normalized median-fluorescent intensity loading at 33 μM for 1 hour (dashed line) with flow cytometry. Data showed mean ± s.d. n = 3–4 independent samples.

We next investigated the relationship between depoted Ag on APCs and loading parameters (i.e. loading concentration and time). A lipid-conjugated long peptide, L-GP100 (CAVGALEGPRNQDWLGVPRQL), was used to prevent direct peptide epitope binding to surface MHCs, requiring internalization before loading and MHC I-restricted presentation [41]. Bone marrow-derived dendritic cells (BMDCs) were generated as indicated in *Methods*, and depoted for up to 1 h with increasing concentrations of L-GP100. Increasing the depot time from 15 min to 1 h resulted in a 2.9-fold increase in L-GP100 at 33 μM, and a 4.3-fold increase at 3.3 μM (**Figure 2E**). The increases were more subtle at 0.033 and 0.33 μM. When depot time was kept constant at 1 h, increasing concentration from 0.033 to 33 μM resulted in a 47.8-fold increase in L-GP100 loading. The loading in BMDCs plateaued after 1 h. We also used L-GP100 to test Ag loading for B-APC delivery. Increasing the depot time from 15 min to 1 h resulted in a 11.7-fold, 9.2-fold, and 5.1-fold increase in L-GP100 at 33 μM, 3.3 μM, and 0.33 μM, respectively (**Figure 2F**). The increase was subtle at 0.033 μM. At a 1 h depot time, increasing concentration from 0.033 to 33 μM resulted in a 173-fold increase in L-GP100 on B-APCs.

To confirm that loaded L-GP100 was processed and presented by B-APCs, we co-cultured loaded APCs with GP100-specific CD8^+^ T cells (PMELs) to assay MHC I-restricted Ag presentation. The highest L-GP100 loading conditions (33 μM for 1 h) were used to load BMDCs and B-APCs prior to co-culture with PMELs for 3 days. BMDCs processed and presented GP100 to cognate PMELs regardless of lipid conjugation (**Figure S2** *left*). In contrast, B-APCs were unable to process GP100 peptide and activate PMELs, yet L-GP100 was processed by B-APCs and presented to activate PMELs (**Figure S2**, *right*). These data suggest that lipid conjugation can facilitate internalization of non-minimal peptide Ags into poorly endocytic APCs for MHC I-restricted presentation.

Ag delivery by L-Ags is accomplished *ex vivo* due to their potential for high levels of non-specific cellular interactions, but this process may be limited *in vivo* due to the susceptibility of DSPE-PEG for hydrolysis [35]. We previously developed polymer-based NPs as an *in vivo* drug delivery system, where encephalitogenic peptide Ags were conjugated to PLGA (P-Ag) and stoichiometrically incorporated into PLGA NPs to induce Ag-specific immune tolerance in a mouse model of multiple sclerosis [24]. Notably, the biodistribution of the PLGA NPs was primarily found in the liver and spleen, where P-Ag could be processed and presented to autoreactive CD4^+^ T cells. We hypothesized that L-Ags along with P-Ags could be incorporated into PLGA NPs to combine advantages of both delivery platforms while overcoming their disadvantages. PLGA NPs can encapsulate hydrophobic polymer conjugates as well as protect encapsulated L-Ags or P-Ags from enzymatic degradation, combining multiple advantages of two drug delivery mechanisms: i) depoting by lipids and ii) endocytosis by NPs to achieve controlled delivery of Ags to B cells and DCs. To comprehensively understand the distinct differences in the drug delivery features between L-Ag and P-Ag NPs, we first conjugated OVA peptide Ags to PLGA using carbodiimide chemistry as controls (**Figure S3**). ^1^H-NMR determined the coupling efficiencies of PLGA-SIINFEKL (P-1; 91%) and PLGA-OVA_323-339_ (P-2; 68.3%). Pairwise combinations of L-Ags and/or P-Ags (**Table 1 and Figure 3A**) were incorporated in PLGA NPs using the single emulsion-solvent evaporation method to create a total of 4 NPs with sizes ranging from ~700-800 nm, zeta-potentials of ~-30 mV, and Ag loadings of 8 μg/mg PLGA NP that was shown in our previous work to sufficiently induce Ag-specific T cell responses (**Table 2**) [24]. These experiments confirmed that L/P-Ag NPs could be prepared with well-controlled physicochemical properties through systematic combination of L-Ag and/or P-Ag conjugates with unmodified PLGA polymer.

**Figure 3.**
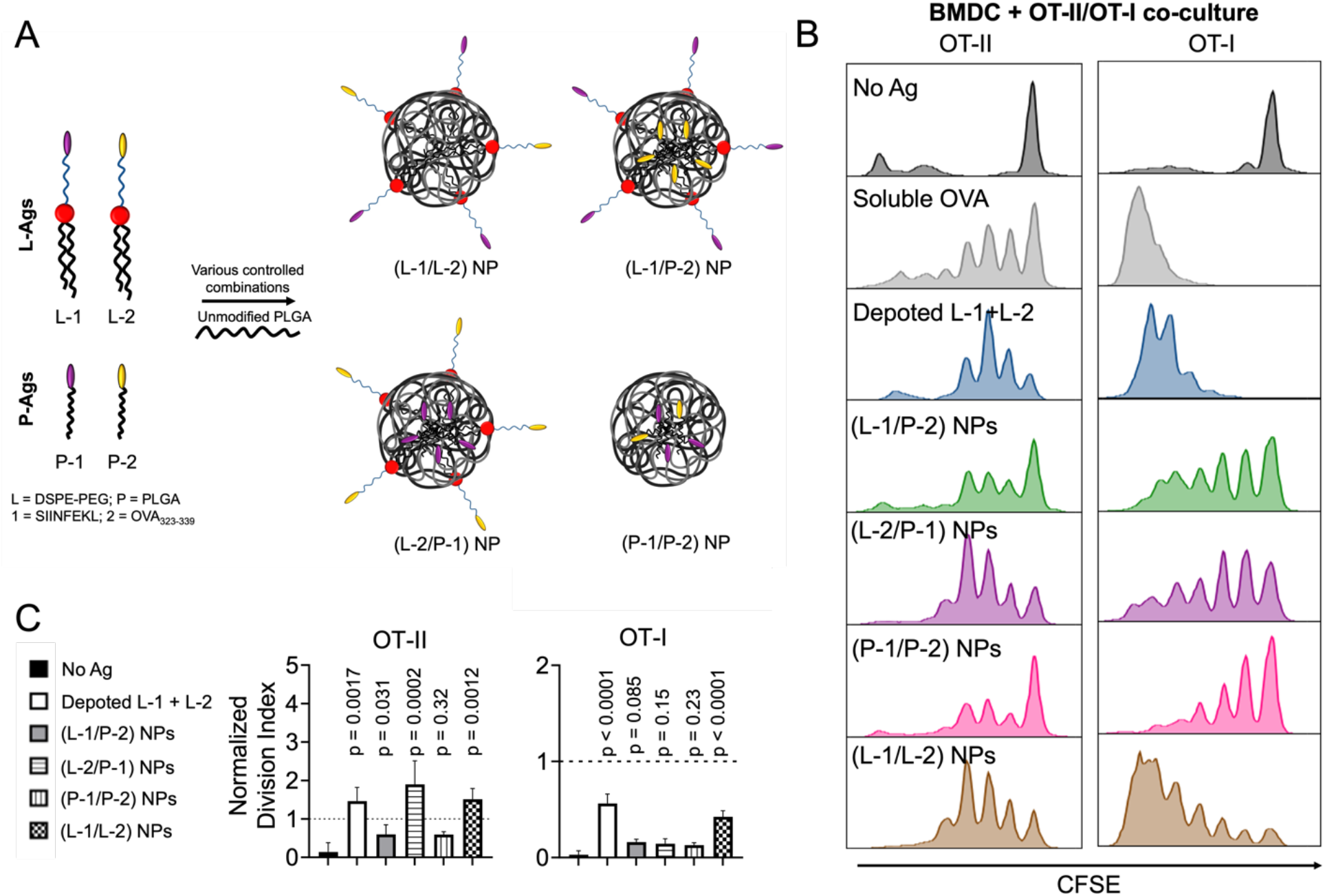
Lipid and PLGA Ag conjugates in NPs enable MHC II- and MHC I-restricted presentation by BMDCs for priming OT-II and OT-I T cells. **A)** Schematic of pairwise combinations of SIINFEKL (1) and OVA_323-339_ (2) Ags with either lipid or PLGA biomaterials, which are then formulated in PLGA NPs. **B-C)** BMDCs were loaded with unmodified OVA peptide Ags, lipid Ag conjugates, or conjugate combinations in NPs: L-1/L-2, L-1/P-2, L-2/P-1, or P-1/P-2. Loaded BMDCs were co-cultured with OT-II or OT-I cells for 3 days. **B)** Representative histograms show T cell proliferation as determined by CFSE dye dilution using flow cytometry. **C)** OT-II and OT-I division indices were normalized to soluble OVA controls, as represented by dashed line. Data showed mean ± s.d. n = 3 replicates. p values determined by using RM one-way ANOVA with Dunnett’s comparisons with no Ag control.

To evaluate the differences in Ag-specific T cell responses induced by various NP formulations, we performed an *in vitro* co-culture experiment. NP formulations were first added to BMDCs followed by the addition of transgenic OVA-specific CD4^+^ T cells (OT-II) or transgenic OVA-specific CD8^+^ T cells (OT-I) and cultured for 3 days. OT-II and OT-I T cells proliferated with all Ag formulations when compared to BMDCs alone (**Figure 3A**). We quantified T cell expansion by calculating proliferation and division indices and normalized these to soluble SIINFEKL and OVA_323-339_ Ags. Depoted (L-1+L-2), loaded (L-1/P-2) NPs, (L-2/P-1) NPs, and (L-1/L-2) NPs increased OT-II T cell normalized division indices by 10.5-fold, 4.3-fold, 13.7-fold, and 10.9-fold, respectively when compared to no Ag control (**Figure 3B**, *left*). Depoted L-1+L-2, loaded (L-1/P-2) NPs, and (L-1/L-2) NPs increased OT-I T cell division indices by 19.3-fold, 5.6-fold, and 14.6-fold, respectively when compared to no Ag control (**Figure 3B**, *right*). Only (P-1/P-2) NPs did not increase T cell proliferation indices (**Figure S4**), suggesting that polymer conjugates do not efficiently promote Ag cross-presentation on MHC-I. Altogether, L-1 and L-2, either with NP loading or without NPs via depoting, delivered Ag to APCs and promoted both MHC class I- and II-restricted presentation, resulting in Ag-specific T cell expansion, while P-Ags in NPs were less efficiently presented on MHC class I.

Activating B cells through CD40 co-stimulation can enhance Ag uptake and presentation by B-APCs, and has been used to prepare B cell-based therapeutic cancer vaccines [7][42][43]. We tested if combining depoting with NPs could deliver L-Ag to CD40 B-APCs. B cells were stimulated with CD40 ligand and TLR7 ligand R848 for 2 days to prepare CD40 B-APCs as previously described [44]. CD40 B-APCs had increased expression of MHC II and CD40, two surrogate markers for cell activation, by 15.2-fold and 3.2-fold, respectively, compared to unstimulated B cells (**Figure 4A**) [16]. Ag loading in CD40 B-APCs was similar to BMDC loading. OT-II and OT-I T cells were co-cultured with CD40 B-APCs at a 1:1:1 ratio for 3 days, then analyzed for proliferation using flow cytometry (**Figure 4B**). Depoted L-1+L-2 increased the OT-II T cell normalized division index by 9.7-fold and normalized proliferation index by 1.5-fold compared to no Ag (**Figure 4C**, *left*; **Figure S5**, *left*). Pulsed OVA did not increase OT-II T cell normalized division index but did increase normalized proliferation index by 1.4-fold (**Figure 4C**, *left*; **Figure S5**, *left*). Depoted L-1+L-2 increased OT-I T cell normalized division index by 35.3-fold and normalized proliferation index by 2.8-fold, while pulsed OVA increased OT-I T cell normalized division index by 34.1-fold and normalized proliferation index by 2.5-fold (**Figure 4C**, *right*; **Figure S5**, *right*). Loaded (L-2/P-1) NPs increased OT-II T cell normalized division index by 6.3-fold and normalized proliferation index by 1.4-fold (**Figure 4C**, *left*; **Figure S5**, *left*). (L-2/L-1) NPs increased OT-II T cell normalized division index by 6.5-fold and normalized proliferation index by 1.4-fold. (L-1/P-2) NPs did not increase OT-I T cell normalized division index but did increase normalized proliferation index by 1.6-fold (**Figure 4C**, *right*; **Figure S5**, *right*). (L-2/L-1) NPs increased OT-I T cell normalized division index by 22.8-fold and normalized proliferation index by 2.2-fold. PLGA-conjugated Ags in NPs were unable to be endocytosed and presented by CD40 B-APCs, similar to B-APCs. These data suggest that CD40 B-APCs can uptake and process L-Ags, either as conjugates or in NPs, and activate cognate Ag-specific T cells as shown by increased T cell responses.

**Figure 4.**
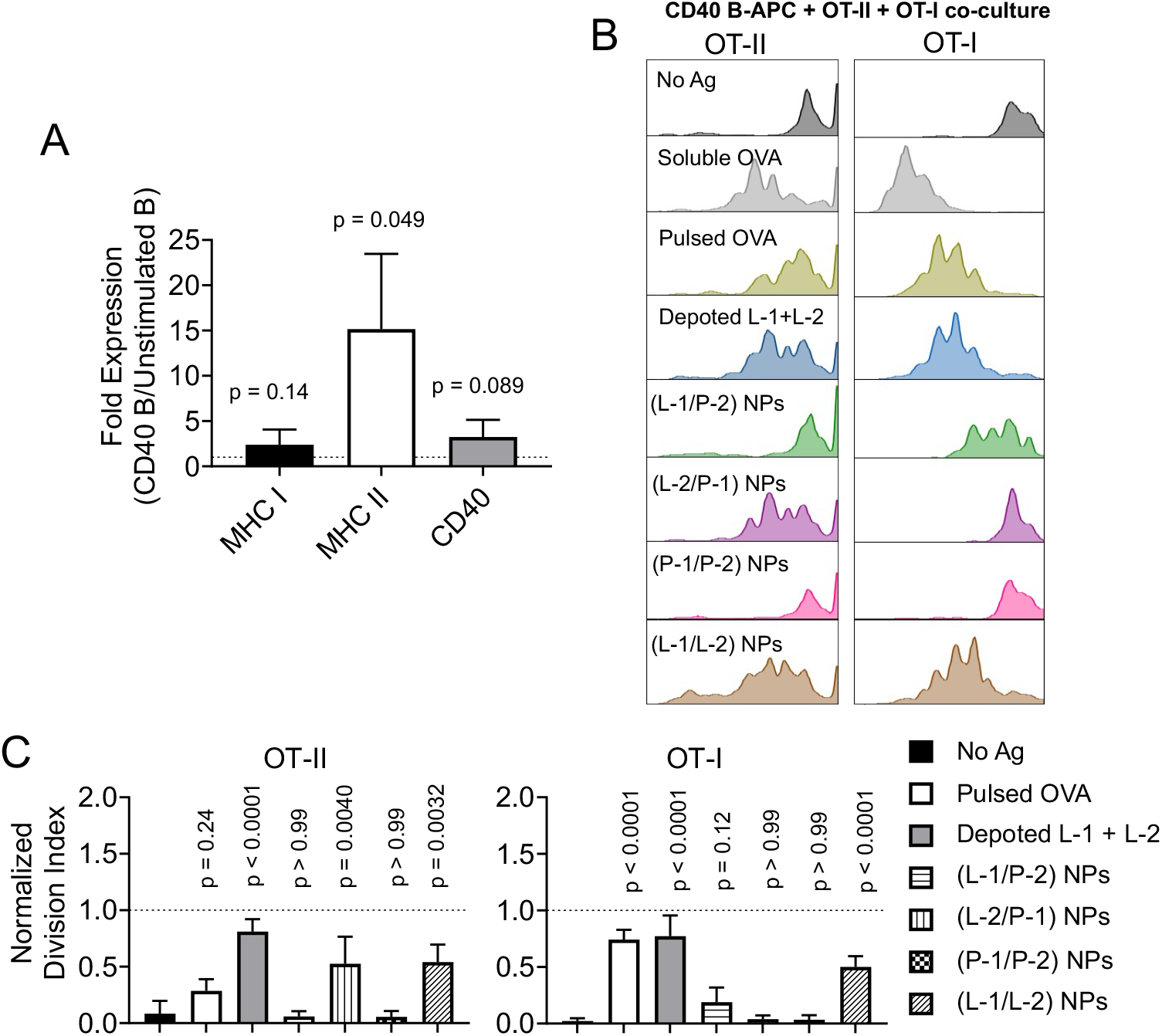
Lipid and PLGA Ag conjugates in NPs enable MHC II- and MHC I-restricted presentation by CD40 B-APCs for priming OT-II and OT-I T cells. Splenic B cells were activated with CD40 mAb and R848 agonist for 2 days. **A)** MHC I, MHC II, and CD40 activation marker expressions were determined by ratios between CD40 B and resting B cell expressions (MFI) using flow cytometry. p values as determined by one-tailed one sample t tests, by comparing to a theoretical fold expression of 1 (dashed line). Data showed mean ± s.d. n = 3 independent samples. **B)** CD40 B-APCs were loaded with unmodified OVA peptide Ags, lipid Ag conjugates, or conjugate combinations in NPs: L-1/P-2, L-2/P-1, P-1/P-2, or L-1/L-2. CD40 B-APCs were then co-cultured with OT-II or OT-I cells for 3 days. Representative histograms show T cell proliferation as determined by CFSE dye dilution using flow cytometry. **C)** OT-II and OT-I division indices were normalized to soluble OVA controls, as represented by dashed line. Data showed mean ± s.d. n = 3 independent samples. p values determined by using RM one-way ANOVA with Dunnett’s comparisons with no Ag control.

Resting B cells are known to be inefficient at non-specific Ag uptake compared to DCs [11]. We next tested if our NPs could effectively deliver Ag to resting B cells (B-APCs) despite lack of CD40-mediated activation. B-APCs were treated with similar groups as CD40 B-APCs, and were co-cultured with OT-II and OT-I T cells at a 1:1:1 ratio for 3 days followed by analysis of proliferation by flow cytometry (**Figure 5A**). Depoted L-1+L-2 increased OT-II T cell normalized division index by 5.0-fold and normalized proliferation index by 1.5-fold when compared to the no Ag control (**Figure 5B**, *left*; **Figure S6**, *left*). However, pulsed OVA peptide Ag did not increase the OT-II T cell division index but did increase normalized proliferation index by 1.4-fold (**Figure 5B**, *left*; **Figure S6**, *left*). Depoted L-1+L-2 increased OT-I T cell normalized division index by 19.7-fold, while pulsed OVA increased normalized division index by 16.1-fold (**Figure 5B**, *right*). Interestingly, OT-I T cell normalized proliferation indices were not increased by pulsed OVA or depoted L-1+L-2 (**Figure S6**, *right*). NPs did not increase OT-II T cell normalized division indices (**Figure 5B**, *left*), but loaded (L-2/P-1) NPs and (L-1/L-2) NPs increased OT-II normalized T cell proliferation index by 1.5-fold each (**Figure S6**, *left*). (P-2/L-1) NPs increased OT-I T cell normalized division index by 6.9-fold, but did not increase normalized proliferation index (**Figure 5B**, *right*; **Figure S6**, *right*). Loaded (L-1/L-2) NPs increased OT-I T cell normalized division index by 20.2-fold, and normalized proliferation index by 2.6-fold (**Figure 5B**, *right*; **Figure S6**, *right*). PLGA-conjugated Ags (P-1 and P-2) in PLGA NPs were unable to induce Ag-specific CD4^+^ or CD8^+^ T cell responses in co-cultures using B-APCs. Taken together, these data suggest that L-Ags delivered either as free conjugates or within NPs can be successfully delivered to B-APCs to activate cognate T cells as shown by increased T cell division or proliferation indices.

**Figure 5.**
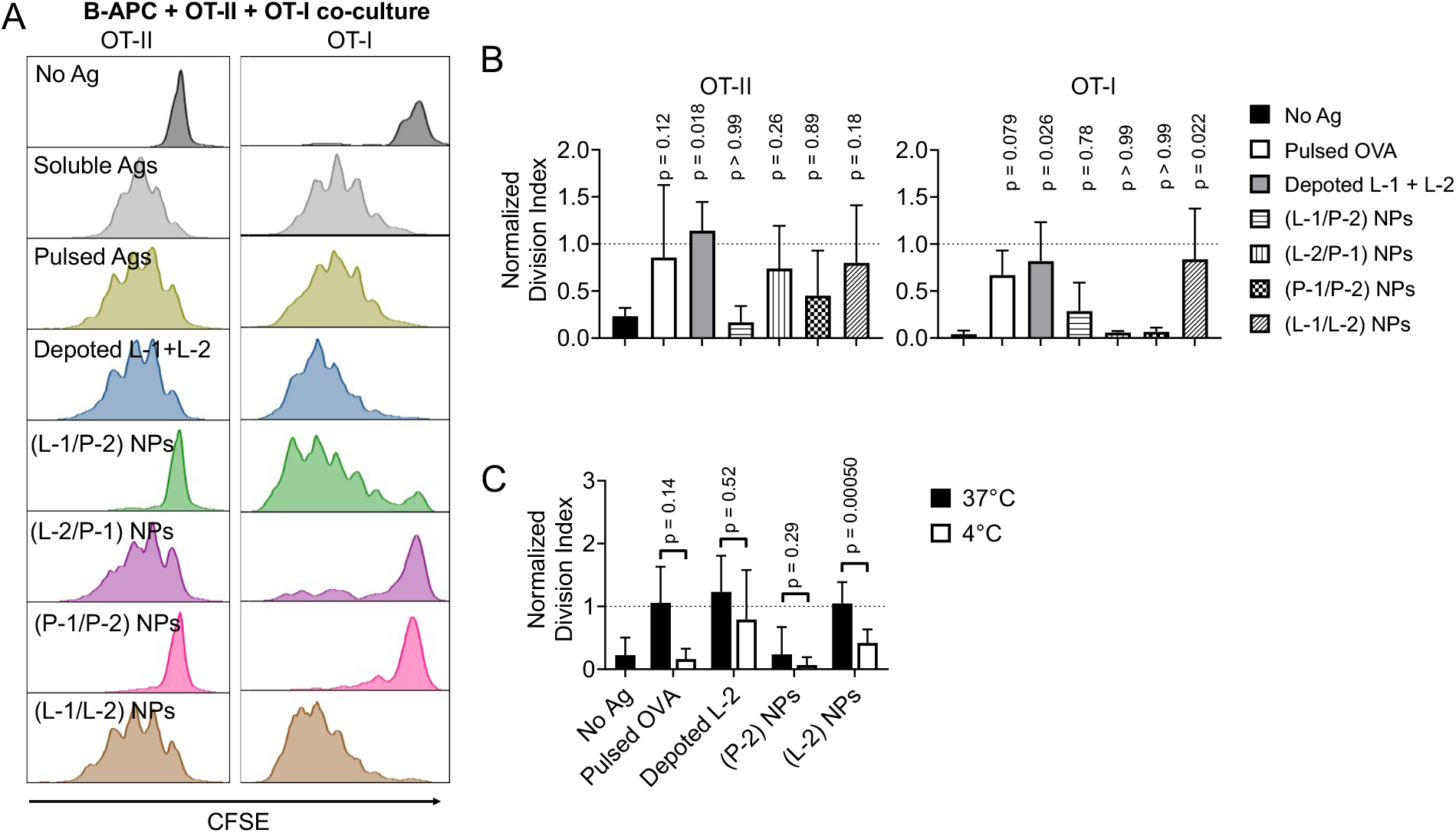
Lipid and PLGA Ag conjugates in NPs enable MHC II- and MHC I-restricted presentation by B cells for priming OT-II and OT-I T cells. B cells were loaded with unmodified OVA peptide Ags, lipid Ag conjugates, or conjugate in NPs: L-1/P-2, L-2/P-1, P-1/P-2, or L-1/L-2. B cells (B-APCs) were then co-cultured with OT-II or OT-I cells for 3 days. **A)** Representative histograms show T cell proliferation as determined by CFSE dye dilution using flow cytometry. **B)** OT-II and OT-I division indices were normalized to soluble OVA controls, as represented by dashed line. Data showed mean ± s.d. n = 3 independent samples. p values determined by using RM one-way ANOVA with Dunnett’s comparisons with no Ag control. **C)** B-APCs were loaded with OVA Ags as lipid conjugates or as lipid/PLGA conjugates in NPs for 1 hour at either 4°C or 37°C, and co-cultured with OT-II cells for 3 days. Division indices were determined by flow cytometry analysis and normalized to soluble OVA control. Data showed mean ± s.d. n = 3–8 independent samples. p values between 4°C and 37°C Ag loading as determined by two-tailed paired *t*-test.

Since B-APCs can process and present L-Ag to induce Ag-specific T cell responses independent of NP-mediated delivery, we tested whether the two delivery methods were mechanistically distinct. We hypothesized that NPs require active endocytosis by APCs for Ag uptake, and this is a temperature-sensitive cellular process [34]. In contrast, L-Ag depoting is passive and less dependent on loading temperature. B-APCs were treated with L-2 or L-2-containing NPs at either 37°C or 4°C for 1 hour before co-culture with OT-II T cells for 3 days followed by analysis using flow cytometry (**Figure 5C**). Depoting L-2 at 4°C did not significantly decrease OT-II T cell normalized division index, but (L-2) NPs loaded in B-APCs at 4°C had a 1.7-fold decrease in OT-II T cell normalized division index. As anticipated, B-APCs loaded with (P-2) NPs and co-cultured with OT-II T cells led to no increase in normalized division index for either temperature compared to the non-stimulated T cell control as similarly observed in **Figure 5B**, *left*. These data suggest that PLGA NP delivery to B-APCs is partially dependent on active endocytic pathways, and delivery of L-Ags in NPs can circumvent a dependence on endocytic pathways and instead leverage delivery via depoting. Thus, depoting offers control over Ag delivery to B-APCs through an expanded mechanism of delivery not available to unconjugated Ags or traditional PLGA NPs.

We next investigated how APC type and Ag formulation affected T cell phenotypes through measurement of cytokine secretions. APCs were treated with various L/P-Ag NPs or relevant controls and subsequently cocultured with Ag-specific OT-II and OT-I T cells for 3 days. Cytokine secretion profiles can be used to broadly define phenotypes, such as CD4^+^ helper or CD8^+^ cytotoxic T cell subsets [45]. Secreted cytokines were quantified using a Luminex-based assay and hierarchical clustering was used to analyze secretory phenotypes of different APC and Ag formulation combinations. No Ag control was distinct from all Ag-containing conditions (**Figure 6A**). OVA pulsed and lipid-mediated depoting were similar, priming T cells that secreted broad, strong inflammatory cytokine milieus. NP formulations had a distinct secretory phenotype from no NP conditions, with L-1 or P-1 formulations subclustering together. This pattern suggested that CD8^+^ cytokine responses were primary drivers of secretory differences between NP formulations. (P-2/L-1) NPs and (L-2/L-1) NPs were most similar, characterized by lack of IL-5 in DC co-cultures and higher MIP-1α and MIP-1β in B-APC cultures. (L-2/P-1) NP and (P-1/P-2) NPs were distinct from L-1-containing NPs. (L-2/P-1) was characterized by high IL-10 in CD40 B-APC cultures, high IL-7 and MIP-2 in B-APC cultures, and G-CSF in DCs. Within the co-cultures containing DCs, IL-2 and TNFα levels were secreted to a lesser extent when OVA_323-339_ Ag was conjugated to PLGA instead of lipid. Ags conjugated to PLGA only (P-1/P-2) NP versus lipid only (L-1/L-2) NP led to increased IL-4 and IL-5 in DC co-cultures, suggesting that delivery of PLGA-conjugated Ags may skew T cell phenotypes towards a Th2 response. Within the CD40 B-APCs co-cultures, IFNγ, IL-2, MIP-1α, and MIP-1β levels increased when Ags were conjugated to lipid instead of PLGA. For resting B-APCs co-cultures, increased IFNγ, IL-1α, IL-2, IL-10, MIP-1α, and MIP-1β levels were observed when Ags were conjugated to lipid and depoted. Notably, IL-7 levels were the lowest when both Ags were conjugated to lipid or PLGA and the highest levels were measured in the (L-2/P-1) NP group. These results demonstrated that the method of Ag delivery plays a distinct role in modulating T cell responses, which offers an opportunity to pair specific APC types and Ag delivery strategies to tune T cell effector and helper phenotypes.

**Figure 6.**
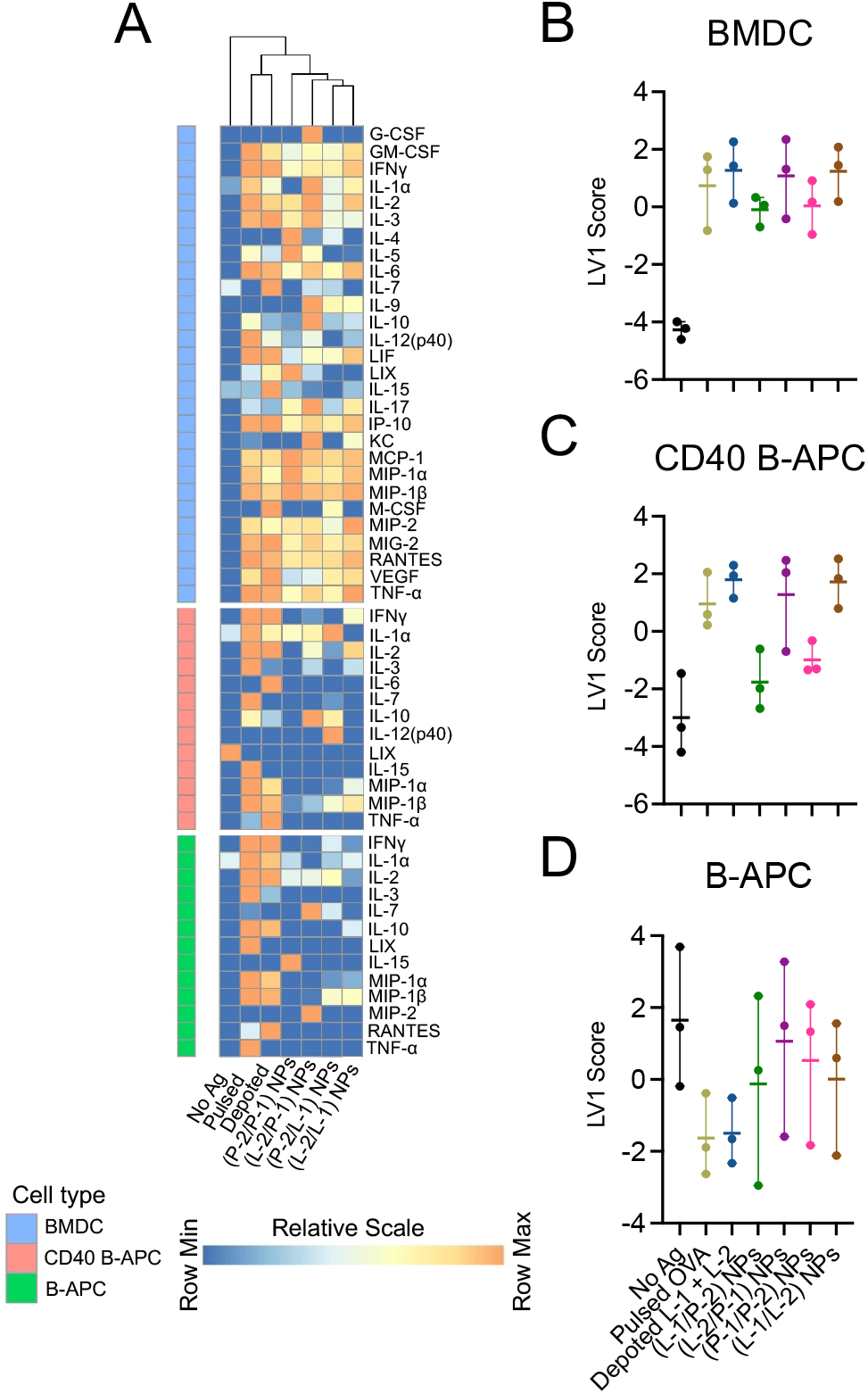
Ag delivery strategy drives cytokine secretion. BMDCs, CD40 B-APCs, and B-APCs were loaded with unmodified OVA peptide Ags, lipid Ag conjugates, or conjugate combinations in NPs: L-1/P-2, L-2/P-1, P-1/P-2, or L-1/L-2. APCs were then co-cultured with OT-I and OT-II cells for 3 days. **A)** Heatmap represents row-normalized concentrations of cytokines for each biomaterial-based Ag treatment as measured by Luminex. Non-secreted cytokines were omitted. n = 2 replicates. **B-D)** PLS-DA model classified scores from latent variable 1 (LV1) from 8 cytokines, TNFα, IFNγ, IL-1α, IL-2, IL7, IL-10, MIP-1α, and MIP-1β, and were grouped based on Ag delivery formulation. Data showed individual points with horizontal line representing mean. n = 3 independent samples.

Utilizing 8 differentially expressed cytokines (MIP-1β, IL-2, TNFα, MIP-1α, IFNγ, IL-7, IL-1α, and IL-10, a multivariate partial least square discriminant analysis (PLS-DA) model was trained to classify different Ag/APC combinations, and variable importance in projections scores were calculated to define significant cytokines on latent variable 1 (LV1). BMDCs separated Ag formulations from no-Ag control due to TNFα, IFNγ, IL-2, and MIP-1α (**Figure 6B**; **Figure S7A**). CD40 B-APCs separated P-2 conjugates in NPs and no-Ag control from L-2 formulations due to MIP-1β, IL-2, TNFα, MIP-1α, and IFNγ (**Figure 6C**; **Figure S7B**). Resting B-APCs separated depoted L-Ags were from no-Ag control due to IFNγ, MIP-1β, and MIP-1α (**Figure 6D**; **Figure S7C**). Classification of Ag/APC combinations were generally driven by inflammatory cytokines for all cell types. Lipid conjugation of Ag led to a more robust cytokine secretion profile, which correlated with the increased proliferative responses in all three cell types evaluated (**Figure 3; Figure 4; Figure 5**). The differential IL-4 and IL-5 secretory phenotype measured for (P-1/P-2) NPs compared to other NPs provided evidence that the mode of Ag delivery is a controllable parameter to modulate Ag-specific T cell responses.

## Discussion

This study developed and characterized the delivery of Ags by L-Ag depoting and compared the delivery of P-Ag or L-Ag conjugates incorporated into PLGA NPs as an Ag delivery platform (**Figure 1**). We demonstrated successful delivery of minimal and non-minimal peptide Ags to APCs, providing access to several possible delivery mechanisms for MHC-restricted presentation. BMDCs and resting B-APCs maintain respective homeostatic levels of surface MHC I and MHC II [46] that can directly bind minimal epitopes. Conjugates consisting of a lipid tail domain and an MHC-restricted minimal peptide Ag may conformationally change to directly bind surface MHCs, or be cleaved by extracellular lipases to liberate Ag epitopes. Additional studies are warranted to discern mechanisms of minimal epitope binding to MHCs. These findings can have important implications in immunotherapies, where minimal Ags that directly bind MHCs on APC surfaces have shown to produce short-lived immunity in cancer or Ag-specific immune tolerance [47][48][49][50]. Non-minimal peptides may be internalized into the cytosol via spontaneous lipid flipping across the plasma membrane or through endosomal vesicles for MHC-restricted presentation after enzymatic cleavage (**Figure 1A**). The processed Ag can then be presented on APC surfaces, a feature we verified with L-GP100 (**Figure 2 and Figure S2**). Future studies utilizing lipid-mediated depoting can deliver more therapeutically relevant Ag candidates to APCs *ex vivo*, including proteins, neo-Ag pools, and tumor lysates to promote internalization and MHC-restricted presentation [51]. This approach can also be complexed with PLGA NPs to deliver multiple lipid- and/or PLGA-conjugated Ags to restrict which cell types can process and present the delivered Ags for controlled *in vivo* Ag delivery applications. Collectively, these results demonstrated the versatile approach of using L-Ags, or L/P-Ag conjugates in PLGA NPs to engineer APCs for controlling Ag-specific T cell activation.

Depoting is facilitated by the lipid tail domain partitioning into the hydrophobic lipid bilayer of plasma membranes in aqueous environments [52], enabling a facile *ex vivo* delivery strategy to all cell types. DCs and B-APCs have drastically different non-specific endocytic capabilities [53], but depoting L-Ags can rapidly engineer both cell types. Delivery platforms that are agnostic to cell type have therapeutic potential by exploring diverse and therapeutically relevant APC candidates for Ag and vaccine delivery [54][55]. We demonstrated controlled Ag dosing to APCs by tuning process parameters such as loading duration and Ag concentration, a feature that can have significant implications in immunotherapy design for eliciting either immunogenic or tolerogenic immune responses [21][24]. For example, lower concentrations of Ags have been shown to preferentially induce regulatory or anergic T cells [56][57]. Increasing Ag delivery above this concentration threshold diminishes the regulatory phenotype [58], suggesting Ag dose can be a main design lever for controlling immune phenotype. Clinically relevant chimeric Ag receptors (CARs) and engineered T cell receptors (TCRs) are endowed with specific affinity for their target Ag, and increasing Ag affinity induces biphasic T cell responses [59][60][61]. This suggests that biophysical properties such as binding avidity and Ag density play key roles in overall T cell responses. In CAR signaling, the Ag density threshold for activation is lower than for cell lysis [62], suggesting that balancing CAR T cell activation and effector functions can be tuned through levers such as controlled Ag dose and density display. Lipid-mediated depoting can provide a rapid, facile delivery strategy for controlled Ag dosing, and therefore Ag density, and utilize APCs to screen next-generation engineered TCR and CAR constructs for cell therapy applications in autoimmune diseases and cancers.

While lipid conjugates have already demonstrated enhanced Ag delivery and T cell activation *ex vivo* and *in vivo* [32][63][64][65], incorporating Ag conjugates in PLGA NPs provides an additional delivery feature to sequester Ags to specific organs and desired APCs *in vivo*. The PLGA NPs used in this study with no modifications are negatively charged, and require endocytic pathways to gain entry into APCs [66], limiting NP access to poorly endocytic APCs. We developed NPs to deliver Ags that are tethered to either lipid or PLGA, conjugates that demonstrated distinct mechanisms for Ag processing and presentation. P-Ag containing NPs induced Ag presentation in only DCs (**Figure 3**), while their analogous L-Ag conjugates were also processed and presented by CD40 B-APCs and resting B-APCs to activate Ag-specific CD4^+^ and CD8^+^ T cells (**Figure 4 and Figure 5**). Selectively pairing Ags with either lipid or PLGA biomaterials can leverage delivery mechanisms that are agnostic as well as dependent on cell type. Rational biomaterial selection can also have inherent immune-stimulating or immune-modulating effects. For example, a palmitoyl-based lipid coupled to a peptide antigen can directly upregulate immunity [63][64]. PLGA particles have demonstrated the intrinsic ability to prevent DC maturation and block immune-stimulating signaling pathways [67][68]. Rationally conjugating Ags to either lipids or polymers and encapsulating in NPs can provide a versatile tool to expand delivery mechanisms and control desired T cell immunity or tolerance.

It is notable that after B-APCs were treated with NPs, unbound particles were removed from the B-APCs by non-gradient centrifugation. This process was likely inefficient in separation of suspension cells from dense NPs, suggesting a limitation of the current study. Over the course of the 3-day co-culture, residual NPs may gradually release P-Ags into solution as exact peptides following hydrolysis of P-Ag, enabling direct MHC binding on APCs for OT-I or OT-II T cell activation [24]. However, since P-Ags in NPs did not elicit T cell activation, the observed Ag-specific responses are not likely driven by PLGA hydrolysis-mediated Ag release from residual NPs. Thus, observed T cell activation was restricted to lipid-mediated delivery.

Lipid and PLGA conjugates can robustly deliver Ags to APCs, where Ag processing and elicited T cell responses are typically highly sensitive to differentiation status [5][69]. We demonstrated that lipid conjugates can potentially decouple Ag loading from activation state by delivering Ags to CD40 B-APCs and resting B-APCs. However, CD40 B-APCs and resting B-APCs did not drive congruent CD8^+^ T cell responses. With CD40 B-APCs, L-Ag formulations expanded Ag-specific CD8^+^ T cells as shown by increased normalized proliferation and division indices (**Figure 4C and Figure S5**). With resting B-APCs, L-Ag depoting did not increase normalized proliferation indices, but did increase normalized division indices (**Figure 5B and Figure S6**). One possible explanation for the disparate outcomes between division and proliferation indices is different delivery formulations may not only promote a distinct Ag processing route, but also dictate differential persistence of MHC-restricted Ag presentation. Even though CD8^+^ T cells typically undergo drastic clonal expansion upon initial Ag exposure [70], suboptimal duration of MHC-restricted Ag presentation by (P-1/P-2) NPs may result in no TCR engagement on nonresponsive T cells. Resting B-APCs may be more sensitive to the differential responses driven by conjugate biomaterial than CD40 B-APCs because CD40 B-APCs have greater Ag presenting capabilities, as shown with increased surface expression of MHCs. Multivariate analysis of cytokine secretions suggested that CD40 B-APCs dictated T cell phenotype depending on biomaterial conjugation to OVA_323-339_ Ag, while resting B-APCs did not differentiate no-Ag control from NP delivery (**Figure 6**). Resting B-APCs shifted T cell phenotype upon Ag pulse or depoting, but Ag delivery in NPs did not differentiate phenotypic response from no-Ag control despite T cell expansion with L-Ag in NPs. However, out of the secreted cytokines, inflammatory cytokines were key drivers in T cell phenotype elicited by all three APCs. An unanticipated finding was related to the distinct Th2 cytokine profile (IL-4 and IL-5 secretions) measured in DC co-cultures by (P-1/P-2) NPs (**Figure 6**). Considering that each L/P-Ag NP evaluated consisted of the same base PLGA NP formulation with various L-Ag or P-Ag conjugates incorporated (**Figure 3A**), this result suggested that the PLGA NP carrier was not the driver of the Ag-specific cytokine profile, but rather the method of Ag delivery contributed to the response observed. Continued investigations are needed to understand how biomaterial conjugates and NPs are dictating nuances in Ag-specific cell expansion as well as driving differential phenotypic profiles.

Our data demonstrated Ag loading into diverse APCs using lipid and PLGA biomaterials to control delivery and dosing. Lipid biomaterial and PLGA carriers can be conjugated to a broad repertoire of molecules, including other proteins, DNA, or RNA, enabling more complex immunotherapy formulation. The flexibility of this platform allows for ease of pairing Ags with other immune-modulators, including adjuvants or checkpoint blockade therapies, which have demonstrated therapeutic synergy with cancer vaccines [71]. Delivering autoimmune-relevant Ags with immunosuppressive drugs can prime tolerogenic APCs to induce Ag-specific immune suppression in autoimmune diseases. Future development of this platform can generate a multiplexed approach for rapid manufacturing of immunotherapy to treat cancers and autoimmunity.

## Supporting information

Zhang et al. Supplementary Materials

## Acknowledgments

The authors wish to acknowledge the support of the University of Maryland School of Medicine Center for Innovative Biomedical Resources, Flow Cytometry Core—Baltimore, Maryland for technical assistance. This research was supported by startup funds from the University of Maryland School of Pharmacy (R.M.P.) and the University of Maryland Baltimore County (G.L.S.), Elsa U Pardee Foundation (G.L.S.), UMBC’s Undergraduate Research Awards (E.M.S. and G.S.), UMBC Summer Faculty Fellowship (G.L.S.), a Supplement for Undergraduate Research Experiences (G.L.S. and E.M.S.), the Commercialization & ENTR REsearch (CENTER) Funding Initiative of the Alex Brown Center for Entrepreneurship at UMBC (G.L.S.), the National Institute of General Medical Sciences of the National Institutes of Health under Award Number R35GM142752 awarded to R.M.P., the UMGCC P30 grant under award number P30 CA134274 from the National Cancer Institute, NIH, and National Institutes of Health (NIH) National Center for Advancing Translational Sciences’ (NCATS) Clinical & Translational Science Awards (CTSA) Program Number 1UL1TR003098. B.L.S. was supported by an NIH-NIGMS Initiative for Maximizing Student Development Grant (R25GM55036). E.M.S. was supported in part by the Nathan Schnaper Intern Program in Translational Cancer Research (NIH R25CA186872). G.L.S. has received royalties from SQZ Biotechnologies. M.H.Z., G.L.S, and R.M.P are inventors on a patent application related to the L/P-Ag NP technology described.

