## Supplementary material for "Lipid-polymer hybrid nanoparticles utilize B cells and dendritic cells to elicit distinct antigen-specific CD4^+^ and CD8^+^ T cell responses": Zhang et al. Supplementary Materials

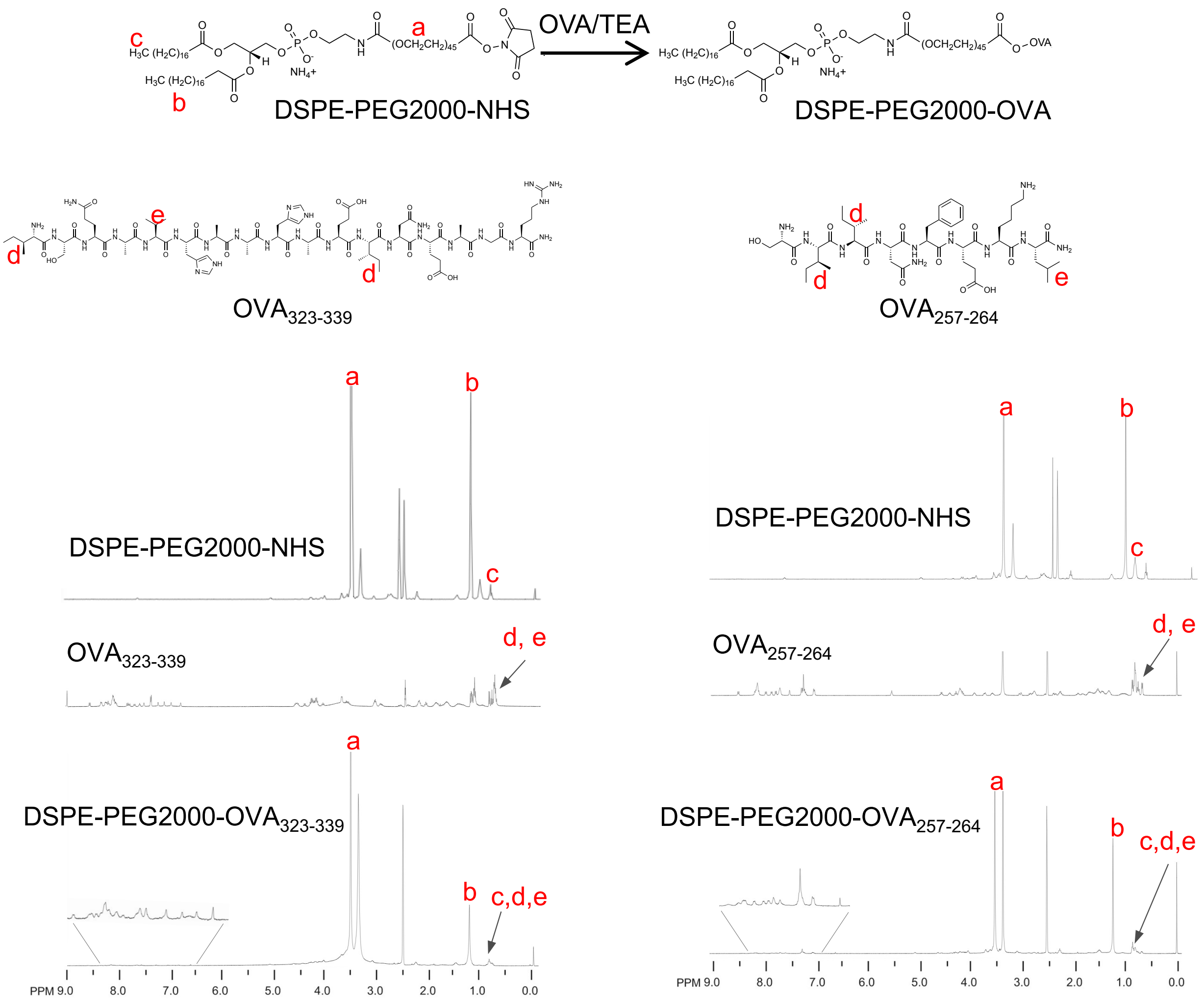


**Figure S1.** Synthesis and characterization of DSPE-PEG2000-$OVA_{323-339}$ and DSPE-PEG2000-$OVA_{257-264}$ (SIINFEKL) conjugates. Measured in DMSO-d6 (calibrated at 2.5 ppm).

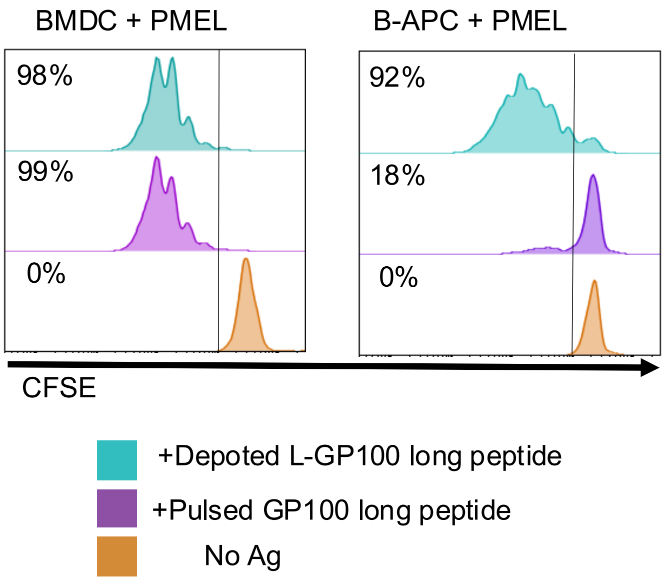


**Figure S2.** Lipid conjugates enables Ag internalization, processing, and MHC I-restricted presentation by APCs for priming cognate and $CD8^+$ T cells. L–GP100 (20-mer) loaded onto BMDCs and B cells was functionally presented to GP100-specific $CD8^+$ T cells (PMELs), as determined by CFSE dye dilution after 3-day co-culture. Percentages represent percent of divided PMELs (as separated by vertical line).

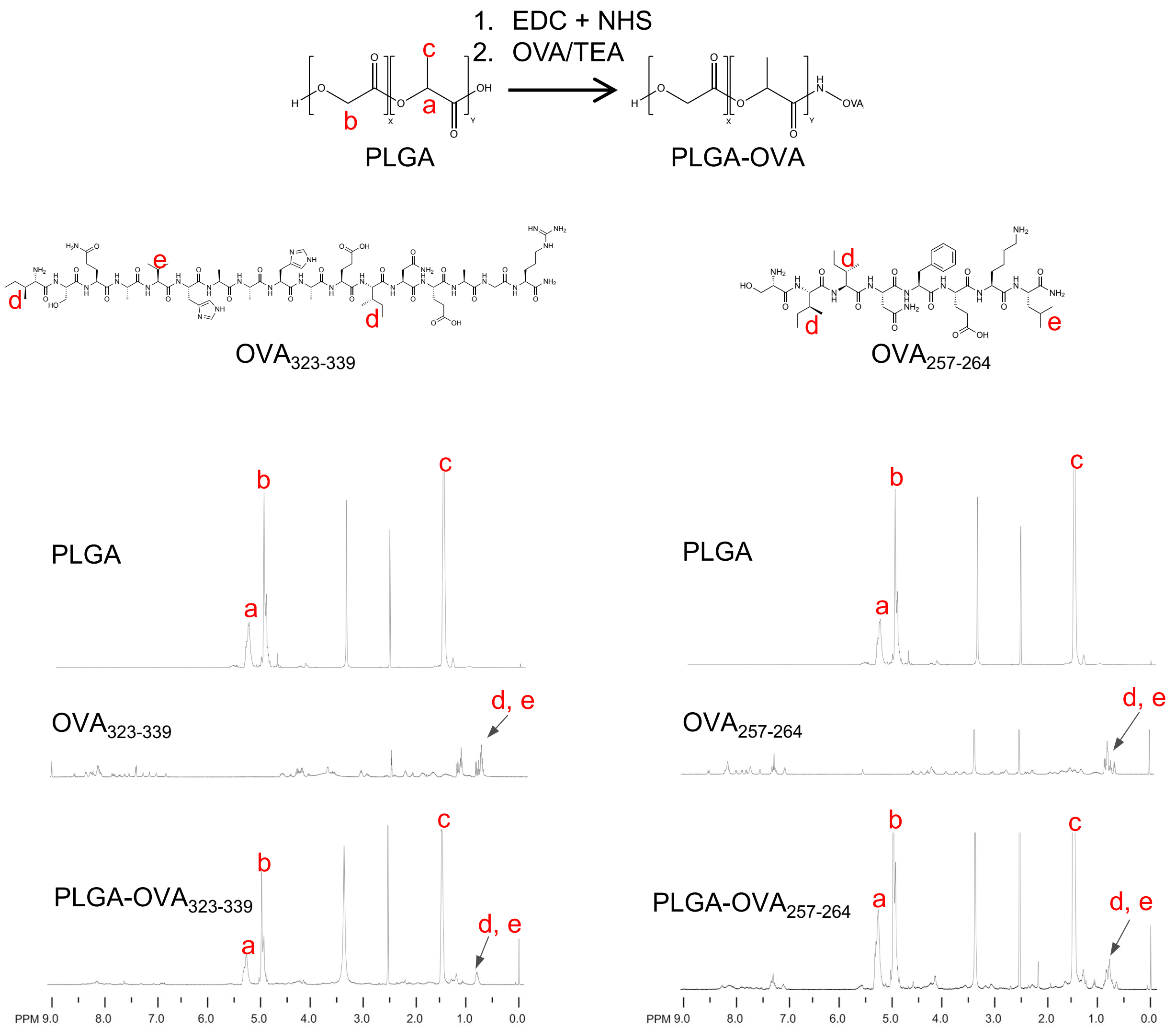


**Figure S3.** Synthesis and characterization of PLGA-$OVA_{323\text{-}339}$ and PLGA-$OVA_{257\text{-}264}$ (SIINFEKL) conjugates. Measured in DMSO-d6 (calibrated at 2.5 ppm).

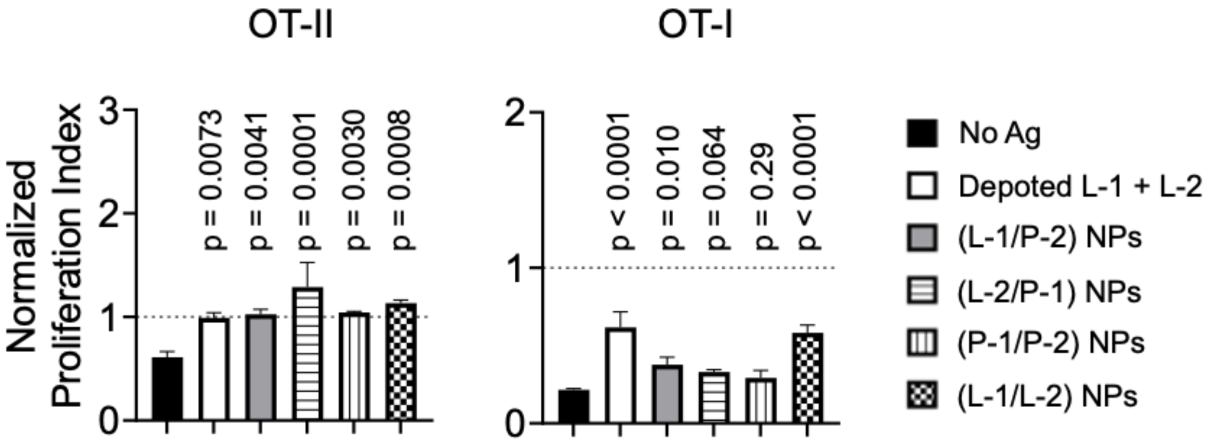


**Figure S4.** Lipid and PLGA Ag conjugates in NPs enable MHC II- and MHC I-restricted presentation by BMDCs for priming cognate CD4$^+$ (OT-II) and CD8$^+$ (OT-I) T cells. BMDCs were loaded with unmodified OVA peptide Ags, lipid Ag conjugates, or conjugate combinations in NPs: P2+L1, L2+P1, P2+P1, or L2+L1. BMDCs were then co-cultured with OT-II or OT-I cells for 3 days. OT-II and OT-I proliferation indices were normalized to soluble OVA controls, as represented by dashed line. Data showed mean ± s.d. n = 3 replicates. p values determined by using RM one-way ANOVA with Dunnett's comparisons with no Ag control.

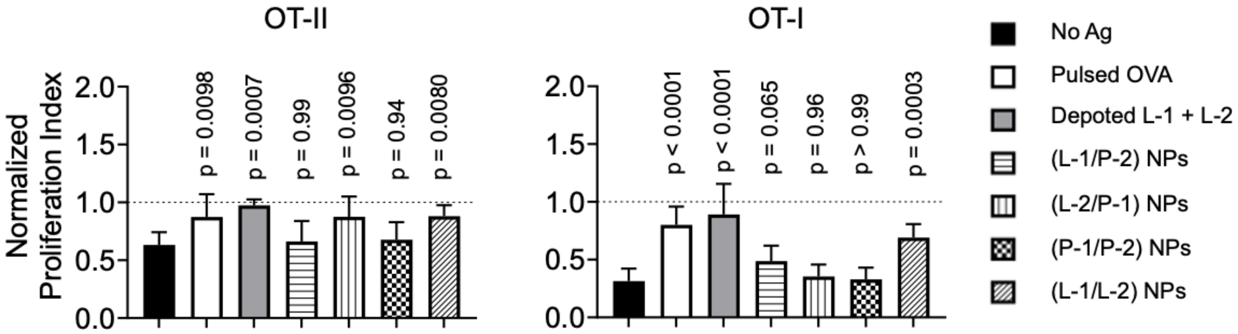


**Figure S5.** Lipid and PLGA Ag conjugates in NPs enable MHC II- and MHC I-restricted presentation by CD40 B-APCs for priming OT-II and OT-I T cells. Splenic B cells were activated with CD40 mAb and R848 agonist for 2 days. CD40 B-APCs were loaded with unmodified OVA peptide Ags, lipid Ag conjugates, or conjugate combinations in NPs: L-1/P-2, L-2/P-1, P-1/P-2, or L-1/L-2. CD40 B-APCs were then co-cultured with OT-II or OT-I cells for 3 days. OT-II and OT-I proliferation indices were normalized to soluble OVA controls, as represented by dashed line. Data showed mean ± s.d. n = 3 independent samples. p values determined by using RM one-way ANOVA with Dunnett's comparisons with no Ag control.

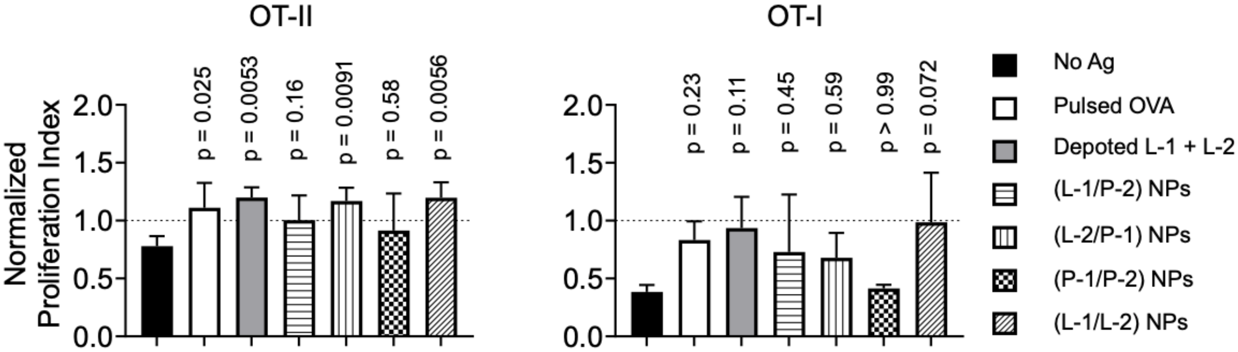


**Figure S6.** Lipid and PLGA Ag conjugates in NPs enable MHC II- and MHC I-restricted presentation by B-APCs for priming OT-II and OT-I T cells. B-APCs were loaded with unmodified OVA peptide Ags, lipid Ag conjugates, or conjugate combinations in NPs: L-1/P-2, L-2/P-1, P-1/P-2, or L-1/L-2. B-APCs were then co-cultured with OT-II or OT-I cells for 3 days. T-cell proliferation was determined by CFSE dye dilution using flow cytometry. OT-II and OT-I proliferation indices were normalized to soluble OVA controls, as represented by dashed line. Data showed mean ± s.d. n = 3 independent samples. p values determined by using RM one-way ANOVA with Dunnett's comparisons with no Ag control.

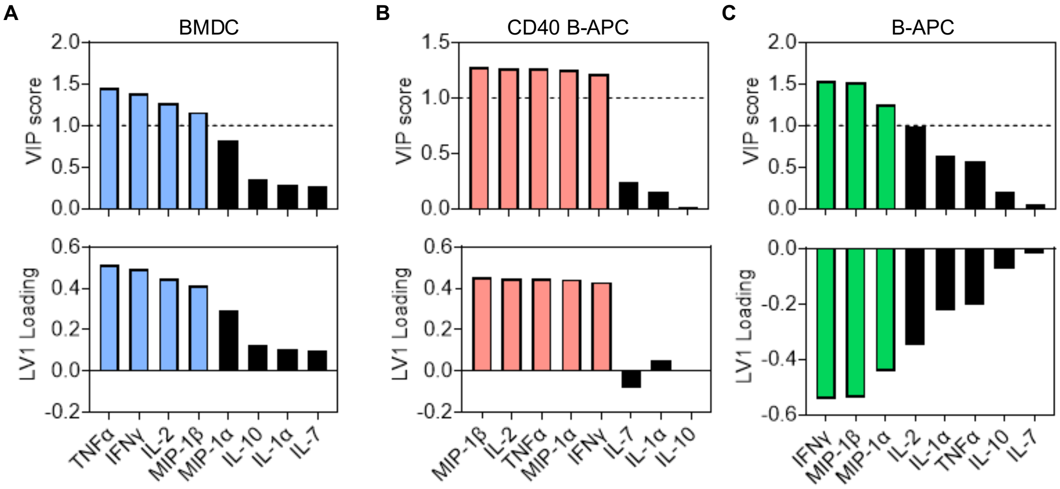


**Figure S7.** Inflammatory cytokines are key contributors to elicited T-cell response. BMDCs, CD40 B-APCs, and B-APCs were loaded with unmodified OVA peptide Ags, lipid Ag conjugates, or conjugate combinations in NPs: P-2/L-1, L-2/P-1, P-2/P-1, or L-2/L-1. APCs were then co-cultured with OT-I and OT-II cells for 3 days. PLS-DA model classified scores from latent variable 1 (LV1) from 8 cytokines, TNFα, IFNγ, IL-1α, IL-2, IL7, IL-10, MIP-1α, and MIP-1β, and were grouped based on Ag delivery formulation. VIP scores > 1 (dashed line) suggested significant contribution to LV1 loading from Ag delivery by **A)** BMDCs, **B)** CD40 B-APCs, or **C)** B-APCs.
